# Comparative genomic insight into the myxobacterial carbohydrate-degrading potential and their ecological impact

**DOI:** 10.1101/2024.11.11.623002

**Authors:** Niharika Saraf, Gaurav Sharma

## Abstract

Myxobacteria are an intriguing group of social-behavior-depicting microbes with unique physiological characteristics such as fruiting body formation, gliding motility, and predation, encompassing the largest genomes (>9 Mb) within the Eubacteria kingdom. These soil-dwelling organisms are crucial for lignocellulosic biomass degradation, which has both ecological and industrial significance. While previous studies have demonstrated polysaccharide deconstruction abilities in a few myxobacterial species, we aim to elucidate the distribution of their Carbohydrate Active Enzymes (CAZymes) domains per organism, with a focus on proteins involved in the catabolism of critical polysaccharides such as cellulose, lignin, xylan, starch, pectin, fructan, chitin, and dextran, across 61 high-quality sequenced myxobacterial genomes. Our findings reveal that 3.5% of the total genes at the median level have domains related to CAZyme functions across different myxobacterial families. Notably, family Archangiaceae (4.4%) and Myxococcaceae (3.7%) members exhibit the most significant genomic diversity and potential for degrading multiple substrates within lignocellulosic biomass. These plentiful CAZymes probably enable these majorly soil-harboring myxobacteria to break down various carbohydrate substrates into simpler biological molecules, which not only allow these organisms to sustain in poor-nutrient environments but also enable them to be critical players in carbon cycling and organic matter decomposition. We conclude that myxobacteria have an unexplored genomic potential that may play an integral role in the degradation of recalcitrant plant biomass, potentially influencing soil health and composition. This study further suggests the critical ecological importance of these CAZymes in sustaining the balance of terrestrial ecosystems and diverse industrial applications.

**Importance:** Polysaccharides are the most abundant polymers making up the Earth’s biomass. Polysaccharide degradation is well-known to be carried out by diverse microorganisms; however, there is more to be explored concerning the novel organisms that can degrade these biomolecules efficiently along with understanding the newer mechanisms and reactions carried out in this process. Soil-dwelling myxobacteria, model organisms for our study, are unique and under-studied social-behavior-depicting microbes. In this research, we investigated their genetic potential to encode carbohydrate-active enzymes involved in breaking down various substrates, including lignocellulosic biomass which is predominantly present in their habitat. We further emphasized their potential to be utilized in industrial applications amongst the paper-pulp, food-beverage, textile, and biofuel industries.

## INTRODUCTION

Carbohydrates, primarily polysaccharides such as cellulose, starch, xylan, chitin, lignin, pectin, fructan, dextran, etc., [1] are the most abundant naturally occurring polymeric biomolecules with numerous imperative biological, pharmacological, and industrial applications [2]. Today, the most abundant biologically renewable resource on earth is the lignocellulosic biomass (LCB) [3], which is primarily composed of cellulose, hemicellulose, and lignin [4]. LCB captures attention as it can serve as an alternative to fossil fuels and pave the way to become the source of biofuels, to meet the ever-increasing demand for energy [5, 6]. Other important polysaccharides are xylan, a component of hemicellulose [7]; chitin, which is abundant in the shells of arthropods like crabs, shrimps, and insects, and produced by bacteria and fungi [2]; pectin, mostly found in plant cell walls and peels of citrus fruits [8]; fructans, which are reserve carbohydrates [9] and dextrans, which are microbial water-soluble exopolysaccharides [10]. The utilization of LCB or other polysaccharides requires physiochemical processing and degradation [11, 12] by diverse Carbohydrate-Active enZymes (CAZymes) [11, 13]. Based on their mechanisms of action, these CAZymes can be distributed into three categories, i.e., 1) enzymes involved in their assembly or glycosyltransferases (GT), and 2) enzymes involved in their breakdown - glycoside hydrolases (GH), polysaccharide lyases (PL), carbohydrate esterases (CE), and 3) accessory functions that contain auxiliary activities (AAs) and carbohydrate-binding modules (CBMs). GHs carry out the hydrolysis of the glycosidic linkage, GTs help in the making of glycosidic bonds, PLs cause the breakdown of glycosidic bonds in a non-hydrolytic manner, CEs hydrolyses the carbohydrate esters, AAs are redox enzymes that play a role in association with other CAZymes, and CBMs help in adhesion to the carbohydrate molecule upon which the enzymes act [14]. Therefore, the understanding of enzymatic reactions behind these processes can help in improving strategies to cope with the recalcitrant nature of some polysaccharides towards degradation [11], further helping in utilizing their potential more efficiently.

Using a diversity of enzymes specific to each carbohydrate substrate, microorganisms degrade miscellaneous carbohydrates in their ecological niches such as the soil [15], oceans [16], or the human gut [17] and maintain equilibrium in those ecosystems. Based on the abundance of these enzymes, LCB decomposition potential has been reported for bacteria, fungi, as well as protozoa [18]. Recent metagenomic studies of rhizosphere microorganisms have shown a plenitude of carbohydrate-active enzymes in Actinobacteria, Proteobacteria, Bacteroidetes, Verrucomicrobia, Gemmatimonadetes and Firmicutes [19]. Bacteroidetes members have also been reported to encode a lot of carbohydrate-degrading genes, playing a critical role in carbohydrate digestion in our gut [17]. The cellulose-degrading potential of *Pseudomonas*, *Streptococcus,* and *Bacillus* species has been widely utilized with several applications in the paper, food, and biofuel industries [20]. However, myxobacteria, which are a notable group of large-genome size non-pathogenic soil-dwelling social microbes, have not been investigated extensively.

Myxobacteria belong to the order Myxococcales within the phylum Proteobacteria, however, recently, the International Committee on Systematics of Prokaryotes (ICSP) classified them under a new phylum, Myxococcota [21, 22]. They are a fascinating group of prokaryotes depicting complex social behavior in the form of fruiting body formation, social motility or swarming, biofilm formation, and predation. Most of these organisms are aerobic with gigantic genomes ranging from 9-16 Mbps, with a few exceptions [23]. Myxobacteria are typically soil bacteria that can thrive in almost any kind of environment, be it temperate regions, tropical rain forests, arctic tundra, deserts, or marine/saline environments. They are known as ‘wolf-pack’ predators and can prey upon bacteria (both, Gram-positive and Gram-negative bacteria) and fungi through toxic secretions into the extracellular environment [24]. Owing to this, they must require a large repertoire of enzymes to degrade lignocellulose biomass and other organisms. An industrially and clinically pertinent characteristic of myxobacteria is that they have a remarkable potential to produce a plethora of secondary metabolites showing a broad spectrum of biological activities such as anti-bacterial, anti-fungal, immunosuppressive effects, and insecticidal, anti-malarial, and herbicidal activities [25, 26]. Overall, these diverse aspects make myxobacteria quite extraordinary and distinct from other bacteria and, therefore, intriguing to study.

Order Myxococcales have been classified into three broad suborders, nine families, 20 genera, and >200 species **(Figure 1, Table S1)** [23]. Most of the members are soil microbes, with a few being marine; however, only some organisms are well studied for their carbohydrate degradation activities, mainly in the context of predation, where these enzymes help in the breakdown of the cell wall of prey bacteria [27, 28]. *Sorangium cellulosum* of the suborder Sorangiineae decomposes cellulose and crystalline cellulose. Most of its strains can also degrade starch, xylan, and chitin [28, 29]. *Sandaracinus amylolyticus*, another Sorangiineae member, degrades starch and has characterized proteins with domains involved in degrading chitin, agar, and cellulose reported in its genome [30]. The Archangiaceae family has several genera among which, *Cystobacter fuscus* and *Stigmatella aurantiaca* are well known to degrade chitin whereas *Archangium gephyra* does not degrade chitin [31]. Chitin degrading activity has also been reported in *Corallococcus sp. EGB* [32] and *Myxococcus fulvus* [33], both of which belong to the suborder Cystobacterineae.

**Figure 1:**
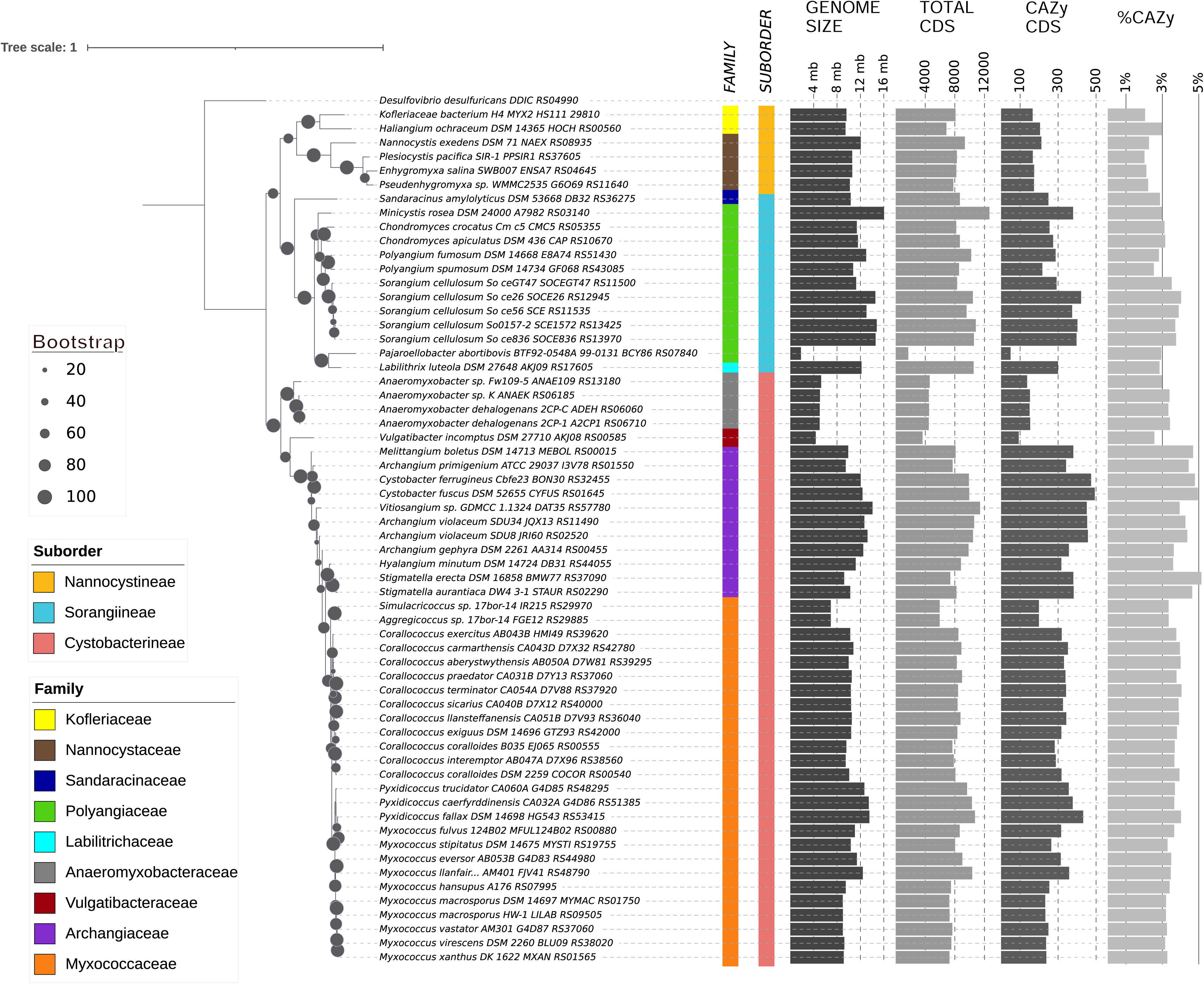
16s rRNA phylogenetic tree depicting the myxobacterial evolutionary relationship at suborder, family, and genus levels along with CAZy distribution: Each label contains the organism’s name with strain and 16S rRNA locus tag. Bootstrap values are represented by the diameter of the circles on the branches of the phylogeny tree according to the scale shown on the left. Taxonomic classification of the three suborders and associated nine families are represented by the color strips according to the left-bottom legends. Bar graphs on the right represent the genome size, CDS counts, CAZy domain containing CDS counts, and percentage of CAZy domain containing CDSs.

Although the carbohydrate degrading potential of a few myxobacterial species has been previously studied, this study aims to perform a high-throughput pan-phylum genomic study to explore the distribution, architecture, and functions of these carbohydrate degrading enzymes (CAZymes). This study investigated sixty-one myxobacterial genomes (majorly complete and a few draft assemblies) and analyzed their CAZymes to untap their polysaccharide-degradation genomic potential. The study highlights varied CAZymes with diverse domain architecture among myxobacterial families, challenging assumptions about their genome size and CAZyme density correlation.

## METHODS

### Selection of organisms

Sixty-one reference and representative myxobacterial organisms were selected from NCBI based on representation of each suborder and family level taxonomy and best possible chromosome-level assembly (draft assemblies were selected only when chromosomal level assembly was not available). Based on assembly and annotation files, a genome statistics table is created to showcase their genomic features such as taxonomy, assembly, NCBI genome ID, genome size, GC%, contigs, CDS number, %CAZy CDS as identified by custom shell scripts **(Figure 1, Table S1)**.

### Phylogenetic analysis using 16s rRNA sequences

16s rRNA sequences of all myxobacterial organisms along with outgroup, *Desulfovibrio desulfuricans,* were extracted and multiple sequence alignment was performed using MUSCLE v3.8.1551. The GTR+GAMMA+I model of RAxML v8.2.12 was used to build the phylogeny tree with 100 bootstraps. Visualization of the resultant tree was done using iTOL v6 (https://itol.embl.de/) [34], along with suborder and family information mapping.

### Domain-based homology detection and domain architecture construction

CAZy domains were identified from the myxobacterial protein sequences by performing hmmscan [35] (HMMER 3.3.2 installed in Nov 2020) against the <u>dbCAN-HMMdb-V9</u> profile database downloaded from http://www.cazy.org/, using default parameters in domtblout output format. The output was parsed using a shell script to sort the output as per overlapped/redundant domains by retaining high-quality matches and calculating the covered fraction of hmm profiles. The parsed and sorted data was represented in the form of ‘domain architecture’ wherein the domains were arranged in order of their position in the protein sequence using a custom R script.

### CAZy distribution

The counts of proteins per organism containing domains specific to each CAZy category were calculated to obtain the distribution values from the architecture. The distribution was obtained at two levels i.e., CAZy broad category level and CAZy family/subfamily level. Custom shell and Python scripts were written to perform these tasks.

### Curation of specific CAZy domains, their architecture and distribution

The different types of CAZymes and their EC numbers were identified through literature search [36–42]. The list of CAZy domains associated with each enzyme was identified by searching respective “EC Number” in CAZYdb. The protein modular architecture specific to those domains was extracted using custom shell scripts. The enzyme-specific domains were also explored to showcase their occurrence in an alone/ duplicate/ combination manner with similar or different domains.

### Correlation and regression analysis

Correlation analyses were carried out using the lm() function in R and the plots for these analyses were made using ggplot2. We performed correlation analysis between genome size and the total number of CAZy domains per organism and similarly between genome size and distribution of each of the six CAZy categories individually.

### Principal component analysis

The principal component analysis was carried out using the prcomp() method and ggplot2 package in R, further clustering using family and suborder information. The principal components used for this analysis were the counts of the six CAZy categories-AA, GH, GT, CBM, PL, and CE per organism.

## RESULTS

### Our studied dataset represents the whole myxobacterial taxonomy

Our dataset included 61 order Myxococcales organisms, which has ∼70% (42/61), ∼10% (6/61), and ∼20% (13/61) representation in three suborders, i.e., Cystobacterineae, Nannocystineae and Sorangiineae respectively **(Figure 1, Table S1)**. We selected these organisms to capture the vast diversity of myxobacteria from all taxonomic lineages with a priority of having a complete genome. As *Myxococcus* and *Corallococcus* spp. from family Myxococcaceae are the most studied and characterized myxobacteria [43], family Myxococcaceae within the suborder Cystobacterineae have the highest representation (26/61; ∼40%) in our dataset. Other predominant families, i.e., Archangiaceae and Polyangiaceae have eleven members each representing ∼20% of the total dataset whereas the rest of the six families have 1-4 members each. Out of twenty-six unique genera, *Myxococcus* and *Corallococcus* have ten and eleven members, respectively. Most of these organisms are soil-dwelling, isolated from diverse lignocellulose biomass present in different countries. However, all suborder Nannocystineae members are present in marine high-salt environments. It is also noteworthy to mention that *Myxococcus fulvus* HW-1 is the only member from the suborder Cystobacterineae that has been isolated from the marine environment [44].

Apart from their habitat and taxonomy, most organisms in the studied dataset have a 9-16 Mb genome size (which is higher than the average bacterial genome size) suggesting their complex metabolic and regulatory implications [45]. *Anaeromyxobacter* spp., *Vulgatibacter incomptus,* and *Pajaroellobacter abortibovis* are exceptions as they have smaller genome size (<5Mb) and a few other physiological characteristics such as respiration, predation, etc., different form the rest of the myxobacteria [46–48]. This dataset includes several largest genomes known so far including *Minicystis rosea* (16.04 Mb genome) [49, 50], *Sorangium* spp. (11-14 Mb genomes) [51], followed by family Archangiaceae and Myxococcaceae members. We found that the myxobacterial family-wise mean genome size is almost 3-4 times greater than the previously reported average bacterial genome size, ∼3.87 Mb [52]. The highest count of total coding sequences (CDS) is present in *M. rosea* and the lowest in *P. abortibovis*. As anticipated, encoded genes per genome show a good correlation with genome size (R^2^ = 0.9648). The GC content ranges from ∼66–75% in our dataset except for *P. abortibovis* with 47.45%. Bacterial GC content can range from 16–75% [53] with *Anaeromyxobacter dehalogenans* 2CP-C at the apex having 74.9% and the other myxobacteria are located in the upper quartile of this range.

### Myxobacteria harbor a diverse CAZy repertoire across their monophyletic taxonomic lineages

First, we employed 16S rRNA sequences of all 61 myxobacterial organisms to build a maximum-likelihood phylogenetic tree to comprehend their evolutionary relationship **(Figure 1)**. This phylogeny was able to perfectly cluster the organisms into distinct clades based on their taxonomic classification at suborder, family, and genus levels. This phylogeny depicts two major clades differentiating into suborder Cystobacterineae members and Nannocystineae and Sorangiineae members together, further separated into their subclades. All families but Archangiaceae show a monophyletic distribution, thereby making a single clade for all members at the genus and species level. However, one of the *Archangium* species within the family Archangiaceae is present at two different branches suggesting the requisite of better phylogenetic markers for their classification.

To understand the relative distribution of CAZy domains and associated polysaccharide degradation capabilities of order Myxococcales, we functionally characterized all myxobacterial protein sequences based on CAZy domains using the HMMER suite’s hmmscan module against the dbCAN CAZy database. We also mapped the organism-wise CAZy CDS distribution to the phylogeny alongside the total CDSs (**Figure 1**) and we saw that some species have a higher CAZyme density. Our analysis reported that myxobacteria utilize almost 2-5% of the total CDSs in carbohydrate-degrading enzyme functions. This study further revealed that members of the suborder Cystobacterineae in the dataset we curated have a slightly higher proportion of CAZy domains in their genomes as compared to suborder Sorangiineae with median values of 3.66 % and 3.11%, respectively **(Figure 2A)**, with the lowest, 2.16%, in suborder Nannocystineae. However, we have more representation from Cystobacterineae members in the present dataset and future studies should include more members from the other suborders. The lower abundance in marine organisms can be explained by their presence in the marine salt-tolerant ecosystem as the meso and bathypelagic layers of the ocean do not have as much availability of organic material for such bacterial species to break down [54]. Family-wise proportions **(Figure 2A)** depict that the highest %CAZy CDSs are present in the family Archangiaceae with a median value of 4.44% and lowest in the family Nannocystaceae with 2.16%. However, more representation is needed from the marine organisms in future analyses.

**Figure 2:**
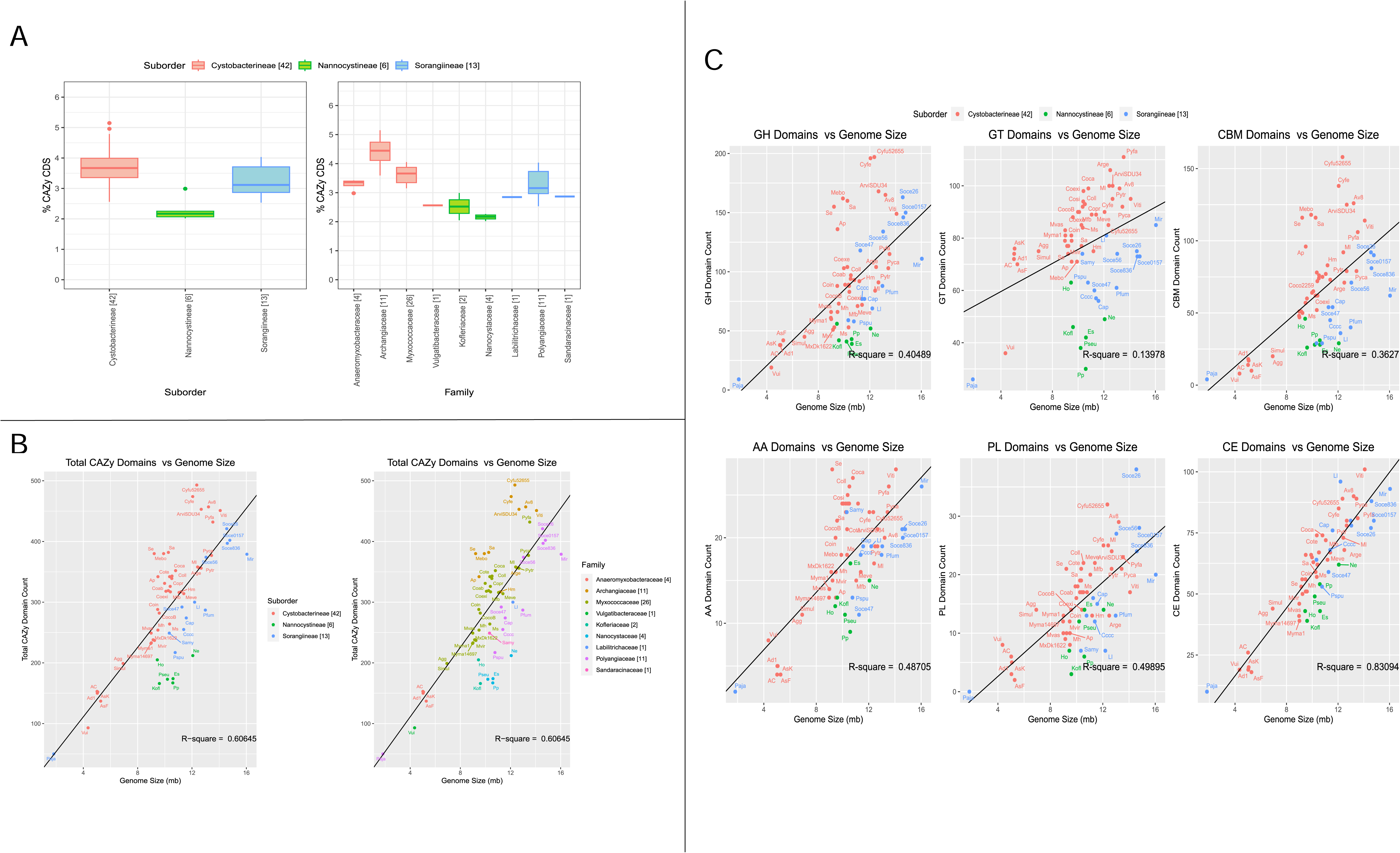
CAZymes distribution and correlation with genome size: **A)** Box whisker plots depict %CDS containing CAZy domains per organism, segregated at suborder level and family level. At the family level, the first four families belong to the suborder Cystobacterineae, the next two families belong to the suborder Nannocystineae, and the last three belong to the suborder Sorangiineae. **B)** Organism-wise correlation analysis between genome size and CAZy distribution counts (the sum of counts of all 6 CAZy Categories-GH, GT, AA, PL, CE, CBM) - showing a) suborder & b) family-wise classification. **C)** Organism-wise correlation analysis between genome size and domain count of each CAZy category (AA, CBM, CE, GH, GT, PL) showing suborder-wise classification. The numbers mentioned in brackets in the legend and the x-axis in all panels depict the number of organisms in that suborder/family.

The overall genomic potential for carbohydrate metabolism can be estimated by counting the number of translated CDSs having at least one CAZy domain from any of the six categories (AA, CBM, CE GH, GT, PL) and we found that the highest number of CAZy domain containing CDSs is 493/9943 **(Table S1)** in the Archangiaceae member *Cystobacter fuscus* DSM 52655 which thrives in soil and herbivorous animals’ dung [29]. We further observed that the five topmost organisms with the highest CAZyme potential have been identified within the family Archangiaceae itself, whereas the lowest number of CAZymes is seen in the Polyangiaceae member, *P. abortibovis*, which is a pathogenic parasite with 1.8 Mb genome size probably having a limited requirement for carbohydrate degradation, and still, it has 2.94% CAZy representation at the gene level. The highest and lowest proportions of CAZymes are present in *Stigmatella erecta* DSM 16858 at 5.14% and *Plesiocystis pacifica* SIR-1 with 2.01% CAZy CDSs in its genome **(Figure 2A)**. Based on this, we hypothesize that CAZyme density is not dependent on the number of CDSs the organism has. To further confirm this, we checked the correlation between genome size and CAZyme proteins **(Figure 2B)**, which revealed that most of the family Archangiaceae members encode more CAZymes as compared to other families. Although we see this potential in Archangiaceae through our comparative genomics approach, experimental validation is required to confirm the expression of these CDSs. Overall, owing to several positive and negative outliers, we found a suboptimal correlation between genome size and CAZy counts (R^2^=0.6). Several organisms deviate from the trendline, and this highlights that despite their relatively smaller genome size, their immense CAZyme potential is an aspect that must be explored and utilized further.

This analysis provided an organism-wise distribution for all 61 organisms for all CAZyme categories, their families, and subfamilies **(Table S2)**. We found that the most abundant CAZyme category is GH followed by GT, CE, CBMs, AA, and PL with median values of 84, 77, 63, 62, 19, and 14 respectively. Archangiaceae members have approximately two times more GHs (160) encoded in their genomes as compared to Polyangiaceae (111) and Myxococcaceae (83), at the median scale. Irrespective of having the largest genome size, *Minicystis rosea,* and other *Sorangium* spp. organisms have almost 30-45% lesser number of GHs, suggesting that Archangiaceae members have the largest genomic potential in terms of carbohydrate degradation as supported by their glycosyl hydrolases encoded proteins. To our surprise, we do not see any major difference in other categories such as GT, PL, CE, and AAs at the median level suggesting their less variation across the dataset. However, CBM numbers showed a large variation of up to two times in Archangiaceae (116) as compared to Polyangiaceae (54) and Myxococcaceae (63), at the median scale, which is almost in correlation with GH numbers. Correlation studies between each CAZyme category with the genome size suggested no correlation for GTs, low correlation for GHs and CBMs (R^2^ value ∼0.38), moderate correlation (R^2^ value ∼0.50) for AA and PL, and good correlation for CEs (R^2^ value 0.83) **(Figure 2C)**.

In the process of seeing an overall picture of the distribution of the major CAZy families and subfamilies across all taxonomic families, we created a ball and dot map of all categories which have an average total of >10 proteins at all taxonomic families **(Figure 3)**. As expected from our above findings, the Archangiaceae family has a greater relative abundance in nearly all major GH families, which are the key domains in polysaccharide degradation. The most abundant subfamilies are CE1 with acetyl xylan esterase activity [55], and other major families include GT4, GT2, GH13, CBM50, and GH23 which are most abundant in either Archangiaceae, Myxococcaceae, or Labilitrichaceae. Among these, relevant to our study are GH13 which has dual roles with α-amylase starch degrading activity as well as glucan-1,6-α-d-glucosidase dextran degrading activity, GH23 which has endochitinase activity, and CE4 involved in acetyl xylan esterase activity.

**Figure 3:**
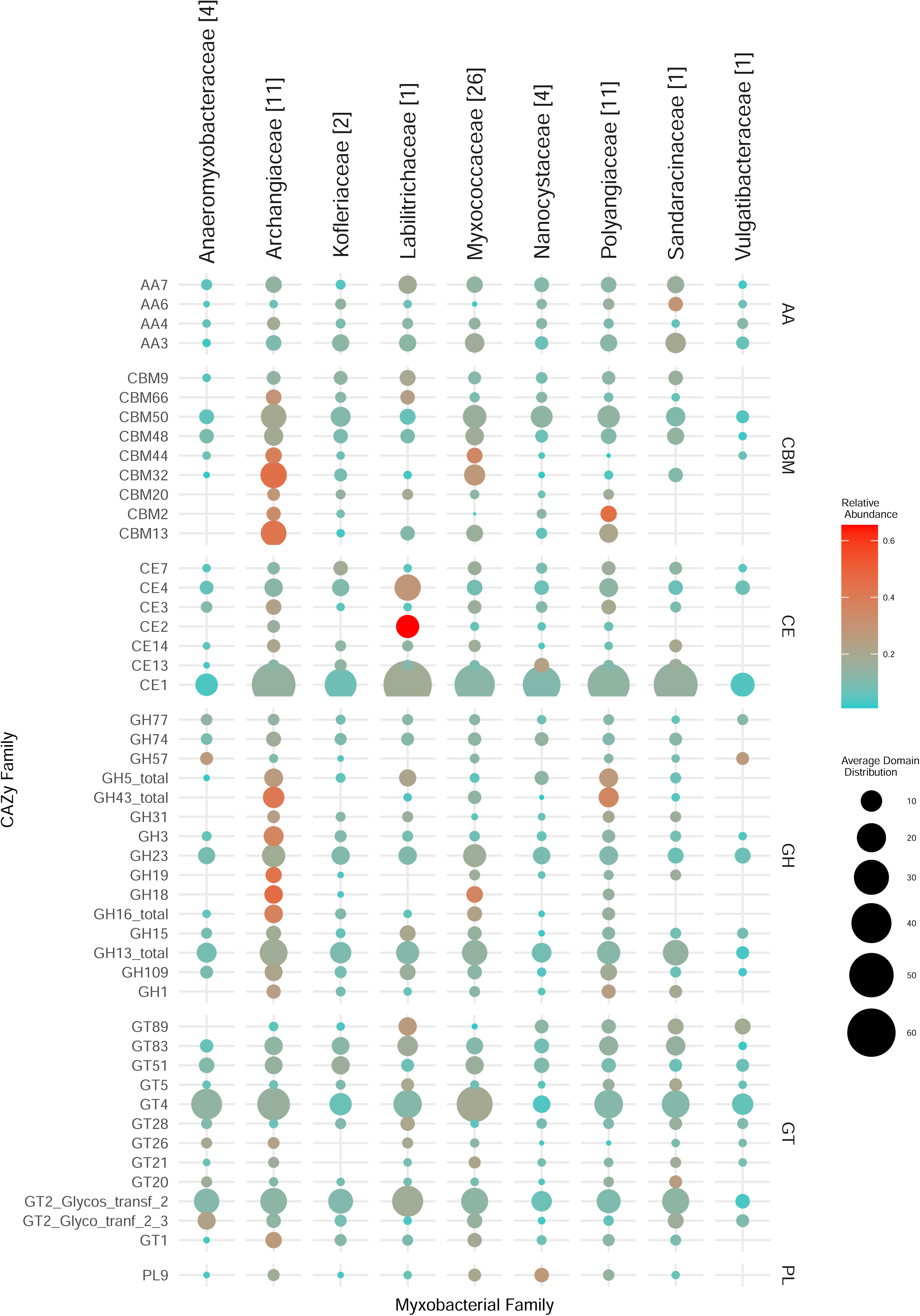
Taxonomy family-wise representation of predominant CAZY families in all six CAZY categories. The mean family-wise values are depicted which are proportional to the diameter of the circles according to the scale shown on the right-bottom. The colour of the circles represents the family-wise relative abundance which gives an idea about how much abundance a myxobacterial family has of a particular CAZy family compared to the rest of the myxobacterial families. Thus, the color range should be analyzed row-wise; blue represents low relative abundance and red represents high relative abundance. The numbers in brackets next to family names (on the top) represent the number of organisms in that family.

We further wanted to analyze if CAZyme density could be used as a factor to differentiate the taxonomic clusters at the suborder and family level and for that we further utilized the distribution counts of the six CAZy categories to perform principal component analysis **(Supplementary** Figure 1). The two axes, PC1 and PC2, showed 72.72% and 10.86% variability, respectively. We observed that at both levels, some clusters were overlapping, justified to some extent by exceptional cases like *P. abortibovis* which have extremely low CAZy counts, and some members of the family Archangiaceae which have relatively high CAZy counts in most categories.

### CAZymes could be playing an important role in the myxobacterial lifestyle

After having looked at the CAZy domain distribution broadly, we emphasized on their functionality based on their actions against major carbohydrate substrates – cellulose, xylan, lignin, pectin, chitin, starch, dextran, and fructan **(Table S3, S4)**. Being majorly soil bacteria, we anticipate that myxobacteria do have a role to play in the degradation of lignocellulosic biomass. Lignocellulosic biomass presents a complex and intricate matrix that is not readily accessible for enzymatic degradation. However, glycoside hydrolases (GHs) and related enzymes exhibit the capability to break down these structures [11], a process that holds significant implications for biofuel production. Consequently, such investigations contribute to the expanding knowledge of these potentially economically valuable enzymes and their natural sources.

Using the enzyme-specific EC numbers (curated from literature) as a query in the CAZy database, we first curated a list of CAZy families that are specific to the enzymatic degradation of each of our chosen substrates **(Table S3)**, followed by their distribution across all myxobacteria **(Figure 4)**. Many CAZy families act on a single carbohydrate substrate but some are bifunctional and involved in the degradation of more than one substrate (the last few domains on the x-axis). For example, GH13 is involved in the degradation of two substrates-starch and dextran. This bifunctionality may be possible because amylopectin (a starch component) and dextran are both α-1,6 linked [56, 57] chains and thus they both can get catabolized by the same enzymatic action i.e. α-glycosidases [58]. Overall, for all enzymes again, the higher abundance is seen in Archangiaceae, Myxococcaceae, and Polyangiaceae members, which needs to be validated experimentally.

**Figure 4:**
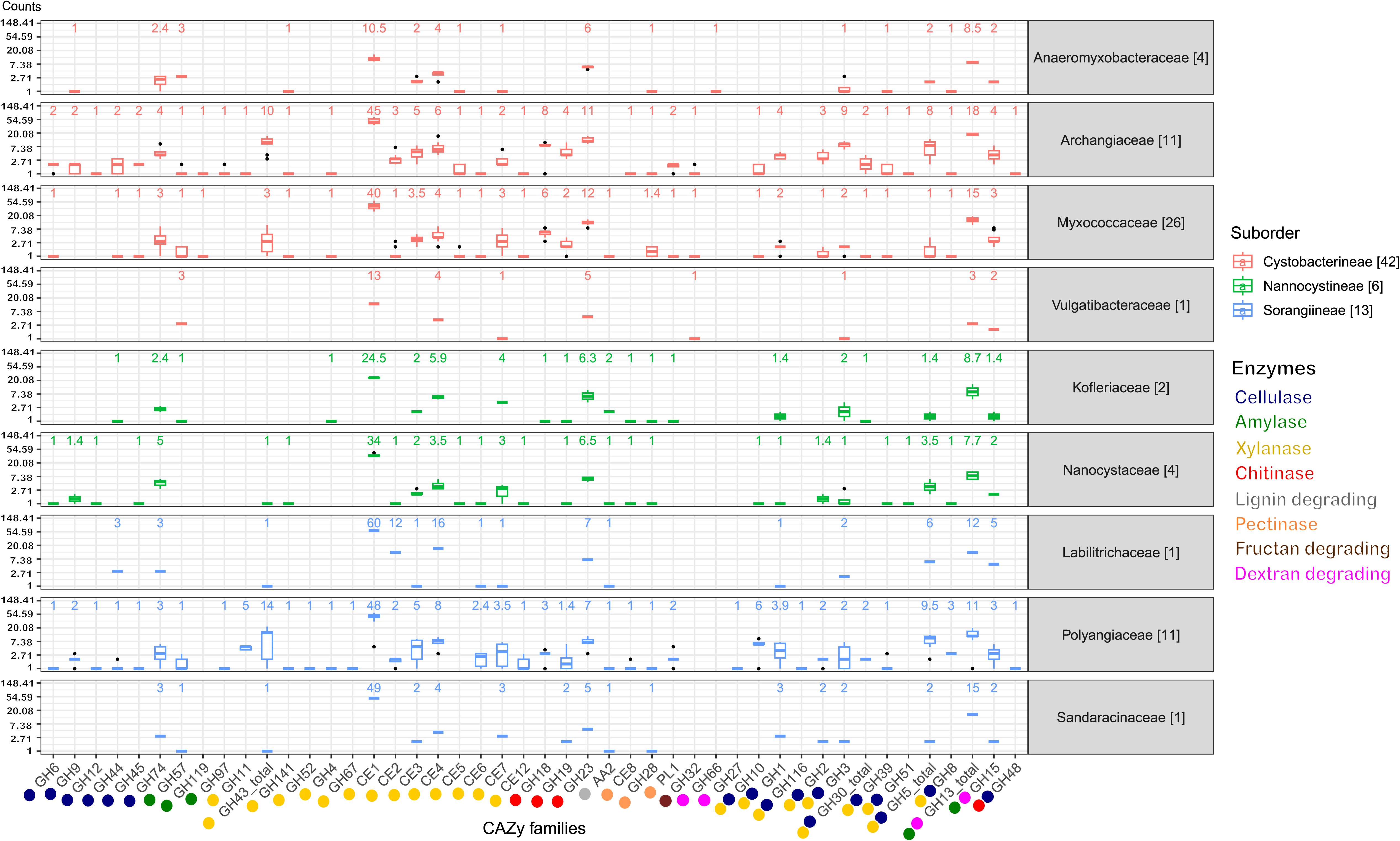
Box whisker plots showing the median count of lignocellulose enzyme-specific CAZy families in the myxobacterial families on a logarithmic scale. The CAZy families are labeled on the x-axis and their corresponding enzymatic function is depicted by the colors below, explained in the lower right corner. CAZy families which are involved in two enzyme activities are represented with both their respective enzyme colours. In the top region of each myxobacterial family’s panel, the mentioned numbers are the median values of that box plot. The box plots are colored according to the suborder. The numbers mentioned in brackets in the legend and beside the family names depict the number of organisms in that family/suborder.

### Myxobacteria are potent lignocellulosic biomass degraders

#### Cellulose degradation

The prime component of lignocellulosic biomass, cellulose is cleaved by different types of cellulases. Seventeen GH families are involved in this catabolic activity, of which six (GH6, GH9, GH12, GH44, GH45, GH74) are unique to cellulase activity in our study, while the others are bifunctional **(Figure 4)**. For 90% of these eleven bifunctional GH domains, the other substrate recognized is xylan while the one remaining, GH48 acts on chitin. The distribution of cellulase domains is highest in the suborder Cystobacterineae owing to the presence of large numbers in a few organisms; however, the median is higher for the suborder Sorangiineae **(Figure 5A)**. The family Archangiaceae harbors 36 cellulase proteins at the median level and Polyangiaceae with 24 cellulase proteins at the median level **(Figure 5B)** suggesting these organisms have the most cellulose degrading genomic potential **(Figure 4, 5B)**.

**Figure 5:**
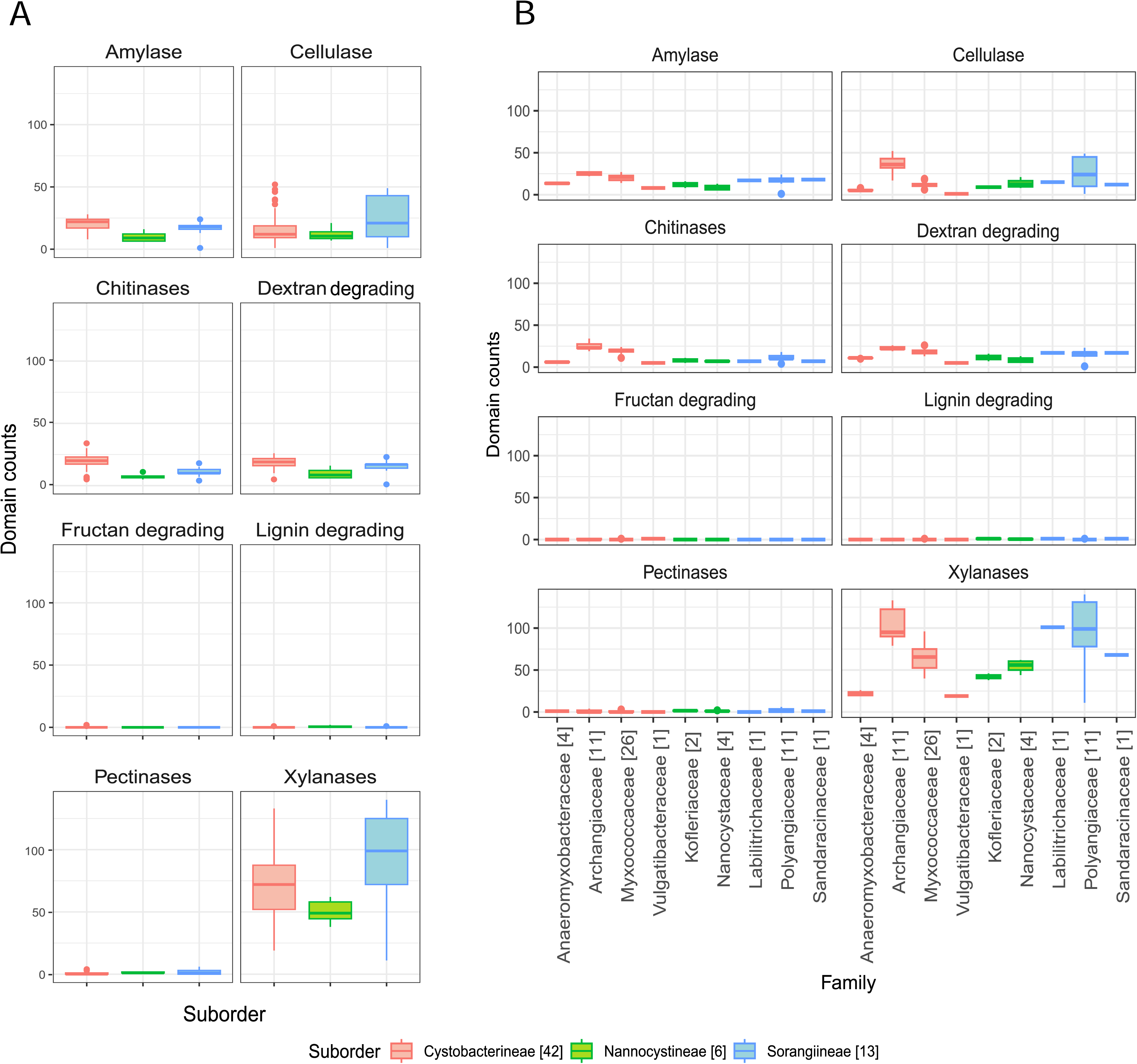
Distribution of specific lignocellulose enzymes: **A)** Enzyme-wise box whisker plots representing the suborder level distribution of the respective CAZy domains. **B)** Enzyme-wise box whisker plots showing the family-level distribution of the respective CAZy domains. The numbers mentioned in brackets in the legend depict the number of organisms in that family. The y-axis scales in all panels are kept the same for comparative visualization.

In Polyangiaceae members, cellulose degradation has already been well-reported [59], both computationally and experimentally. However, within the family Myxococcaceae, cellulases have been reported in a few studies such as encoded by the *cel9-cel48* gene cluster in *Myxobacter* sp. AL-1 [60]. The Polyangiaceae central tendencies in our study, however, are on the lower side due to the exceptional case of the pathogen *P. abortibovis*; the median shoots up to 30 proteins when calculated excluding this pathogen. The members of both these families mostly inhabit the soil ecosystem, in decaying plant material, dung of herbivores, and wood/bark [29] all of which are rich sources of cellulose. Thus, their CAZyme potential reflects their habitat and role in the ecosystem.

The Archangiaceae member *C. ferrugineus* Cbfe23 exhibits the highest number of cellulase-associated domain-containing proteins with a total of 52 **(Table S4)**. Following closely are *S. cellulosum* So ce836 and *S. cellulosum* So ce26, each containing 49 cellulase domain-containing proteins **(Table S4)**. *Vulgatibacter* species from the family Vulgatibacteraceae and *Haliangium ochraceum* SMP-2^T^ (different strain - *Haliangium ochraceum* DSM 14365 used in our study) from Kofleriaceae are known to be non-cellulose degrading [47] This non-degradative characteristic is evident from their lower domain distribution in our dataset as well **(Figure 4, 5B)**.

The cellulase GH domains can occur alone, in duplicates, or paired with other CAZy domains **(Figure 6)**, which possibly means that they can perform their action independently and in association. GH6, GH9, GH48, and GH51 have both endocellulase and exocellulase (or cellobiohydrolase activity) **(Table S3)**. Notably, GH6 is predominantly associated with CBM4, and occasionally with CBM3 and CBM37. Amongst these associated domains, CBM4 is implicated in the degradation of amorphous cellulose and xylan but not crystalline cellulose; CBM3 targets cellulose and, less frequently, chitin; while CBM37 has a broad polysaccharide binding potential, as documented in the CAZy database. GH9 domains are majorly associated with CBM30, CBM4, and CMB3 which are well known to have cellulose, xylan, or chitin-binding function. It is interesting to note that the endocellulase domain GH74 domain is typically present alone in most identified proteins; however, when it co-occurs with other domains, it is linked with GH33, which has trans-sialidase activity. Additionally, GH10, GH1, GH116, GH2, GH3, GH30 (and its sub-families), GH39, GH51, GH5 (and its sub-families), and GH8 are commonly involved in the degradation of both cellulose and xylan.

**Figure 6:**
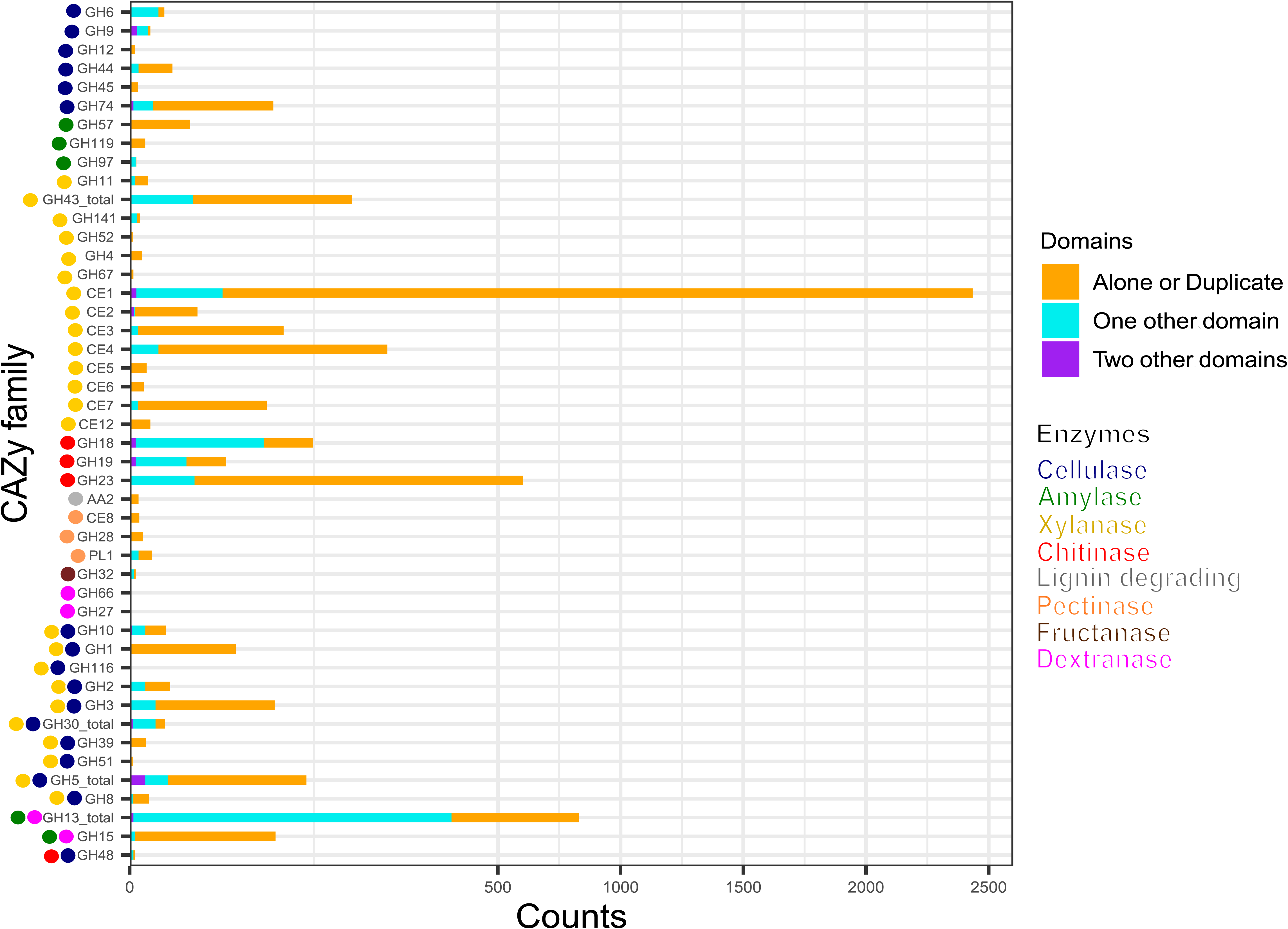
The stacked bar plot representing the relative frequency of CAZy domains in diverse combinatorial patterns. The CAZy domains either occurring alone (or in multiple copies) or with one or multiple accessory domain indicate their independent/ dependent mode of action with one or two functions. The X-axis scale is expanded within 0-500 counts.

We further observed that some cellulase CAZy families were specifically present in a few organisms or families or suborders. GH10 (former cellulase family F), GH12 (former cellulase family H), GH8 (former cellulase family D), and GH9 (former cellulase family E) were predominantly present in genus all under study *Sorangium* spp. and a few members of family Archangiaceae **(Table S3)**. More importantly, these members were almost absent in the most abundant family Myxococcaceae except for sporadic distribution in a few organisms. Amongst the cellobiases (beta-glucosidases) function, we found a unique family, GH116, which is only present in *Sorangium cellulosum* So ce56 (locus_tag=SCE_RS22635); even the taxonomic distribution in CAZy database shows its exceptionally low occurrence in Proteobacteria phylum. Blastp analysis reveals its closest homologs in *Acetivibrio cellulolyticus*, a phylum Bacillota member known for its efficient crystalline cellulose degradation capabilities. Overall, this further suggests the novel origin and functional importance of GH116 protein in *Sorangium cellulosum* So ce56. GH48 (former cellulase family L) and GH51 are also interesting candidates to explore further as they are only present in several family Archangiaceae members and almost absent in other myxobacteria. Overall, cellulase protein distribution indicates the huge cellulose degradation genomic potential within *Sorangium* spp. and less-explored family Archangiaceae members.

#### Lignin degradation

To enhance the fermentation rate of cellulose and hemicellulose, lignin must first be removed which can be performed by bacteria, fungi, bacteria-fungal co-culturing, or enzymatic pre-treatment [61]. Our analysis indicates that auxiliary activities (AA) families AA1 and AA2 are implicated in various ligninolytic processes. Specifically, AA1, which exhibits laccase activity, is absent in all organisms examined in this study. In contrast, AA2, which is associated with lignin peroxidase, manganese peroxidase, and versatile peroxidase activities, appears infrequently and cannot be correlated with their taxonomy. Overall, lignin-degradative domains are not prominently represented in our dataset **(Figures 4, 5)**. AA2 domain does not co-occur with any other CAZy domain **(Figure 6)** suggesting its independent functions.

#### Xylan degradation

Plant hemicellulose predominantly comprises xylan, a polysaccharide structurally similar to cellulose but composed of D-xylose monomers [7]. Cellulose hydrolysis is facilitated by hemicellulose-degrading enzymes, a critical factor for enhancing lignocellulosic biomass degradation in industrial applications [62]. *xynA,* a xylanase-encoding gene has been thoroughly studied in *Sorangium cellulosum* So9733-1 [63]. Xylan degradation is mediated by xylanases, the most diverse class of enzymes in our dataset, spanning 24 different CAZy families. Consequently, there are more xylanase domain-containing proteins per organism compared to those targeting other polysaccharide substrates. Nearly 60% of these CAZy families are specific to xylan deconstruction, with carbohydrate esterases (CEs) being highly represented; no other polysaccharide-degrading enzyme category in our dataset encompasses as many as eight CE families **(Table S3)**.

The function of feruloyl esterases and acetyl xylan esterases is primarily mediated by CE domains. Notably, family CE1, involved in both enzymatic actions, is the most prevalent domain **(Figure 4)**, ranging from five proteins in the Sorangiineae pathogenic member *P. abortibovis* to 64 proteins in the Archangiaceae member *Vitiosangium* sp. GDMCC 1.1324. At the suborder and family levels, significant xylan-degrading potential is observed in Sorangiineae **(Figure 5A)**, particularly within the Polyangiaceae family and Archangiaceae members **(Figures 4, 5B)**. *S. cellulosum* So0157-2 exhibits the highest xylanase distribution with 140 proteins, followed by *S. cellulosum* So ce836 and *S. cellulosum* So ce26 with 138 and 137 proteins, respectively.

Most specific xylan-degrading domains occur independently **(Figure 6)**, though GH43 and its subfamilies, as well as CE1, can possess accessory domains. GH43 subfamilies co-occur with several carbohydrate-binding modules (CBMs): CBM6 which interacts with amorphous cellulose, β-1,4-xylan, and glucan; CBM66, which binds with terminal fructoside residue of fructans; and CBM13, which binds mannose in plants and xylan in bacteria. CE1 domain is occasionally accompanied by a few other CAZy modules such as CBM20, PL22, CE7, and CBM32.

One of the β-xylosidases, GH52 CAZy family, is uniquely distributed only within four out of five *Sorangium* spp. members. These *Sorangium* proteins show maximum homology with *Agarivorans* spp. members from Gammaproteobacteria with 98% query coverage and ∼50% identity. Similarly, the GH67 domain functioning as α-glucuronidase is also present only in *Sorangium* spp. with closest relatives in Gemmatimonadaceae bacterium having >80% query coverage and >60% identity.

### Myxobacteria encode a diverse set of enzymes to degrade other important polysaccharide substrates

#### Starch degradation

At the suborder level, Cystobacterineae members have the highest median values for amylase domain-containing proteins **(Figure 5A)**. Previously, α-amylases, AmyM [64], and CoMA [65] have been characterized in the Cystobacterineae member *Corallococcus* sp. EGB. Another multifunctional amylase, AmyC, has been characterized within the same species [66]. Looking at the family level, Archangiaceae members have the highest median value of 25 proteins **(Figure 4, 5B)**. The Archangiaceae members – *C. fuscus* DSM 52655, *A. violaceum* SDU34, and *A. violaceum* SDU8 have equal numbers and the highest counts with a value of 28 proteins **(Table S4)**. Starch-degrading domains are also found in Myxococcaceae, with a median count value of 21 proteins. The order Sorangiineae and family Sandaracinaceae member, *S. amylolyticus DSM 53668,* is a species known to degrade starch [30] and it has 18 proteins with starch-degrading domains. Starch degradative domains include α-amylases associated GH13, GH57, GH119, β-amylases functioning GH14, γ-amylases associated GH15, GH97. GH13 and its subfamilies as well as GH15 are bifunctional domains with both dextran and starch degrading functions **(Figure 4, 6)**. GH13 subfamiles GH13_11 and GH13_10, were found to be associated with CBM48 in 143 and 102 proteins, respectively which makes it the most frequent accessory domain for GH13 in our dataset. Other accessory domains include CBM20, CBM32, starch binding CBM25 the α-glucans amylose, amylopectin, pullulan binding CBM41, starch binding CBM69 which is also distantly related to CBM20 and CBM48.

GH119 domain is involved in α-amylase function, and it is specifically present in all family Archangiaceae members and only one genus in the family Myxococcaceae, i.e., *Corallococcus* members. GH119 is closely related to the previously well-known α-amylase family, GH57. These myxobacterial GH119 homologs have close homologs in *Cohnella* and *Paenibacillus* spp. of phylum Bacillota (Firmicutes) as indicated by >95% query coverage and >60% identity suggesting their putative evolutionary relationship with the latter.

#### Chitin degradation

Endochitinase activities are predominantly shown by GH18, GH19, GH23, and GH48 CAZy families. A recent study has reported a GH19 enzyme, C25GH19A, which is chitin-induced in *Corallococcus silvisoli* c25j2 [67]. An anti-fungal GH18 chitinase, CcCti1, was also identified and further studied in *Corallococcus sp*. EGB [68]. Based on the distribution of these families, the suborder Cystobacterineae members demonstrate encoding potential for chitin degradation **(Figure 5A)**. Within this suborder, members of the family Archangiaceae exhibit the highest median number of chitinase domain-containing proteins, with a median value of 23 **(Figures 4, 5B)**. *Vitiosangium sp.* GDMCC 1.1324 possesses the highest number of chitin-degrading domains, totalling 34 **(Table S4)**. Among Archangiaceae, previous studies have shown that members of the genus *Cystobacter* can degrade chitin but *Archangium gephyra* cannot [31, 69]. Following the previous study, our study shows that *A. gephyra* DSM 2261 has *a* lower count of chitinase-associated domain-containing proteins as compared to a higher count (19 proteins) in the *Cystobacter* members *C. fuscus* DSM 52655 and *C. ferrugineus* Cbfe23 (29 and 30 proteins respectively). Among Myxococcaceae, chitin-degrading enzyme domains have been previously reported in *Corallococcus* species and *Myxococcus* species [68] [33]. With a median value of 20 proteins, Myxococcaceae members stand second highest after Archangiaceae. Chitinase domains GH18, GH19, and GH23 showcase endochitinase activity amongst which the first two are always majorly associated with other domains whereas the last one majorly works independently **(Figure 6)**. GH18 plays a role in chitobiosidase activity as well and in our data, it is majorly associated with several CBMs, such as CBM12, CBM13, CBM2, CBM44, CBM5, CBM8, CBM13, CBM61, and CBM 37. Similarly, GH19 is present with CBM12, CBM13, CBM2, and CBM50. However, GH23 is mostly independent (515 proteins) but when associated with an accessory domain, it is one or two or three replicates of CBM50 itself. This suggests that CBM50 has some additional evolutionary relatedness with GH23. GH48 acts both on cellulose as well as chitin (endochitinase activity). Our study revealed that this domain is only present in a few members of the family Archangiaceae along with two members of the family Polyangiaceae, i.e., *Minicystis rosea* DSM 24000, *Polyangium fumosum* DSM 14668.

#### Dextran degradation

Suborder Cystobacterineae has the highest genomic potential for dextran degradation as well **(Figure 5A)**. Here again the most potent are the members from the underlying Archangiaceae family **(Figure 4, 5B)** with a median of 22 proteins having dextran degrading domains. *A violaceum* SDU8 has 26 proteins with dextranase domains which is the maximum value in this dataset. GH66 and GH49 play a role in dextranase activity, GH13 with its subfamilies and GH15 are involved in glucan-1,6-α-d-glucosidases functioning and GH27 in glucan-1,6-α-isomaltosidases. GH49 also has a role to play in dextran 1,6-α-isomaltotriosidase activity but GH49 is not identified in any of the genomes in this study.

The GH66 domain is only present in *Anaeromyxobacter* sp. Fw109-5, which depicts the closest homologs in *Thermoanaerobacter* spp. from phylum Bacillota (Firmicutes) with >95% query coverage and ∼45% identity. Both of these are facultative Fe (III)-reducing bacteria, and such homology possibly suggests the shared evolutionary relationship between these two homologs. Similarly, the GH27 domain which is involved in glucan-1,6-α-isomaltosidases function occurs only in *Sorangium cellulosum* So ce836 and not in any other genome. Its closest homologs are present in the *Verrucomicrobiia* bacterium from phylum Verrucomicrobiota, with >70% query coverage, >40% identity, and e-values of <1e^-135^, showcasing good sequence homologs.

#### Fructan degradation

Fructan degradation is catalyzed by fructan exohydrolases which function with the help of the GH32 domain. GH32-containing proteins are not too abundant in this myxobacterial dataset **(Figure 4, 5)**. This domain is only present in six proteins **(Figure 6)**, out of which it occurs independently in two proteins and the other four, with two copies of domain CBM66 which is well-known for fructan binding.

#### Pectin degradation

CE8, GH28, and PL1 are the domain families substantial for pectinase activity CE8 having pectin methylesterase activity, GH28 for endo-polygalacturonases, and PL1 having pectin lyase activity are the domain families substantial for pectinase activity. We could identify only a few pectinases in our dataset **(Figure 4, 5)**, where the highest number is seen in *S. cellulosum* So ce26 with 6 domains. Pectin-degrading pathways have also been earlier reported in *S. cellulosum* [59]. Overall, CE8 was found in only 3 proteins which are encoded by *Archangium violaceum* genomes **(Figure 6)** and our analysis reveals that all 3 of those proteins have varying architecture, one having CE8 in triplicates, one with the domain PL9_4, and the other with the pectinase specific PL1 itself. GH28 is found in 18 proteins overall, while mostly present alone, this domain could also be found associated with one accessory domain which is PL9_2, or two accessory domains PL9_2 and PL9_4 **(Figure 6)**. The accessory PL9_4 plays a role in the peptidoglycan lyase activity, while no specific activity has been characterized for PL9_2 as per the CAZy database. PL1 mostly occurs alone, or it can be with other accessory domains such as PL9, mannose/xylan binding CBM13, the rhamnose binding CBM67, and CBM37.

Overall, our data indicates that the Archangiaceae, Myxococcaceae, and Polyangiaceae families have significant carbohydrate-degrading genomic potential, warranting further exploration, experimental validation, and utilization. Conversely, the pathogenic *Pajaroellobacter* demonstrates the lowest distribution of CAZymes across all classes, which is consistent with expectations for a pathogenic organism that has reduced genome size. For all enzyme categories, the lower values of Nannocystineae could be attributed to their marine habitat [70] (**Figure 7)** and thereby reduced carbohydrate degradation but more representation and experimental validation are required in future studies. This is supported by earlier reports during the initial characterization of *Nannocystis exedens*, which noted the absence of starch and cellulose degradation [71]. Collectively, these findings from the current dataset suggest that CAZyme potential is higher in soil-dwelling myxobacterial species compared to their marine counterparts, though further studies are needed to confirm this trend **(Figure 7)**.

**Figure 7:**
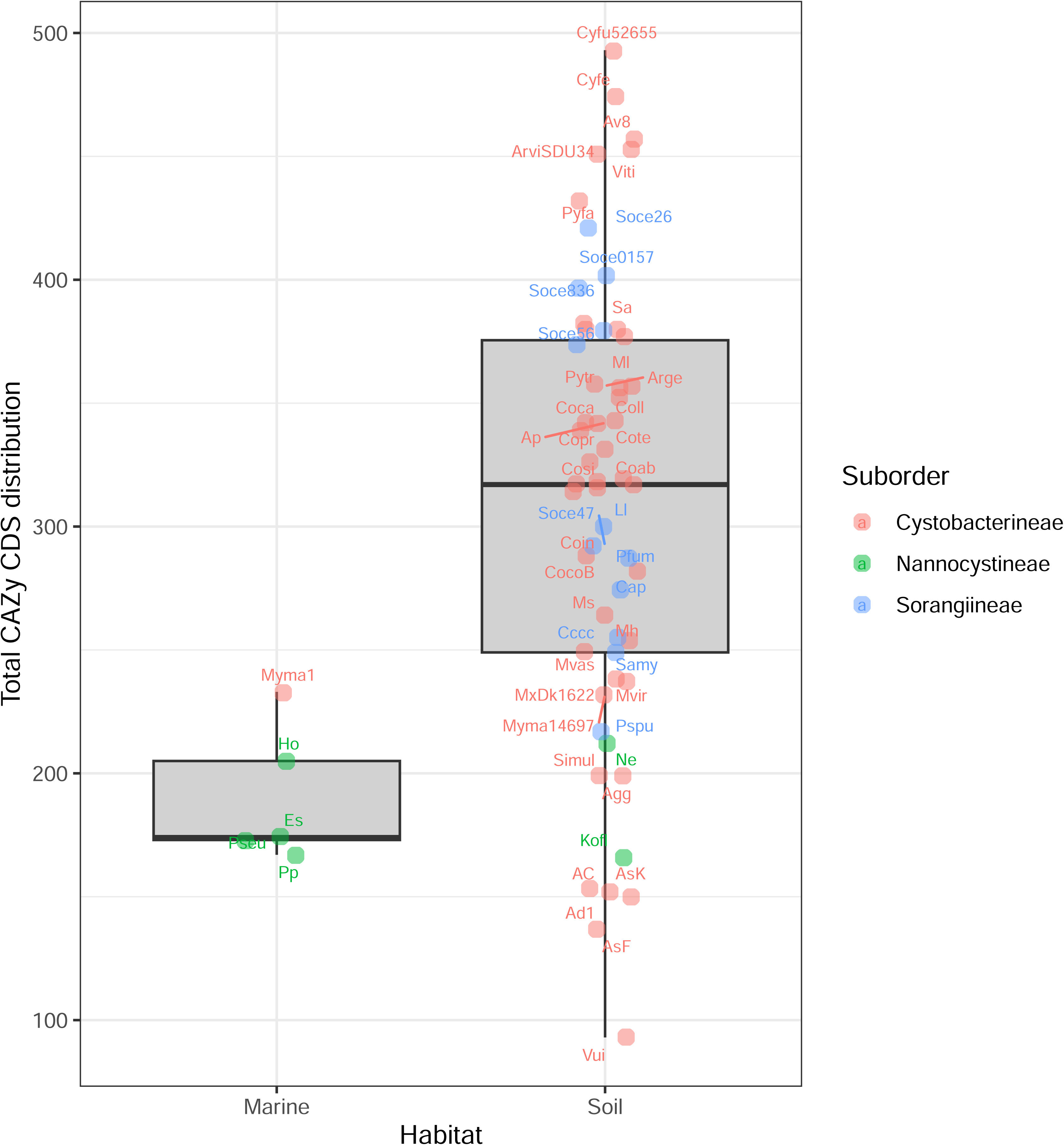
Boxplots showing the organism-wise preferred habitat and respective CAZy domain distribution of the 61 myxobacteria in our dataset, categorized by suborder.

## DISCUSSION

Myxobacteria, ubiquitous across various habitats, exhibit their highest diversity in steppe and warm climatic zones. Notably, certain species within the genera *Myxococcus*, *Corallococcus*, *Archangium*, and *Nannocystis* are also found in harsher environments such as deserts and high-altitude regions. These topsoil bacteria are frequently isolated from the dung of herbivorous animals and decaying plant material, which serve as well-known predominant sources of complex polysaccharides [72]. The genomic potential of organisms in polysaccharide catabolism using the carbohydrate-active enzymes (CAZy) database can underscore their potential for industrial applications.

In this study, we revealed the genomic potential of polysaccharide catabolism in a diverse representation of myxobacterial species by diving deep into analysing the CAZyme domains involved in degrading the major substrates cellulose, starch, xylan, fructan, lignin, pectin, dextran, and chitin. Members of Sorangiineae, especially *Sorangium cellulosum* have been well-known as cellulose degraders as the name suggests, however, the genomic background of these activities was not much studied. Our research fills this gap and reveals that many other myxobacterial species possess considerable, yet underexplored, capacities for polysaccharide deconstruction. The degradation of polysaccharides, especially lignocellulosic biomass, holds immense potential for biofuel production. Previous research has shown that overexpression of certain plant-associated domains, like glycosyltransferases (GTs), can enhance biomass formation [73]. This concept could be explored and applied to harness the converse catabolic activities of myxobacteria for increasing efficiency.

Our findings indicate substantial variability in genomic CAZyme potential among these large-genome harboring myxobacterial families and organisms, with some species exhibiting high potential for substrate-specific degradative capabilities despite having relatively small genomes. Notably, members of the soil-dwelling Archangiaceae family demonstrated a remarkable richness in CAZymes. It is important to recognize and mention that deriving biological conclusions from high-throughput studies such as ours, which involve multi-organism and multi-level comparisons, heavily relies on the quality of publicly available genome assemblies. Most of the assemblies used in our study have complete genomes with high-quality genome annotations, however, a few of them are incomplete, multi-contig assemblies that may not capture the total encoding potential. The inclusion of these draft assemblies was to avail the maximum taxonomic and genomic diversity available at the time of data curation. Despite these limitations, our deep analysis has unveiled new avenues of knowledge.

While we performed this analysis at three levels-suborder, family, and organism, there may be cases where a bacterium/its family may have immense genomic potential but is not expressing the same when it comes to its expression in its life cycle. Genomic studies alone cannot address these limitations, necessitating extensive experimental validation. We urge experimental researchers to verify whether high counts of substrate-degrading CAZy domains correlate with increased degradation efficiency. Nonetheless, the carbohydrate degradation potential highlighted in our study is substantial and provides a basis for designing experiments to harness this capability for practical applications. The dataset used in the study is taxonomically broad but still limited to certain well-studied myxobacterial groups. This could introduce phylogenetic bias, where the observed patterns in CAZyme distribution might not be generalizable to other, less-studied phylogenetically close groups. We acknowledge this limitation and suggest the inclusion of more taxa in future studies to provide a more complete picture.

In conclusion, we present a comprehensive high-throughput genomic analysis of the Carbohydrate Active enZymes-encoding potential of sixty-one myxobacteria members (order Myxococcales or currently known phylum Myxococcota), examining their genomic diversity at the levels of suborder, family, and genus, as well as their functionality in terms of singular or multiple enzymatic activities. Comparative analysis among members of order Myxococcales revealed variations in carbohydrate utilization potential and recommended that lignocellulose biomass degradation is not universally conserved and similar within these groups of organisms. Such carbohydrate degradation analyses lay a preliminary foundation for utilizing myxobacterial organisms in diverse paper-pulp, food-beverage, textile, and biofuel industries.

## Declarations

NA

## Abbreviations

CAZy: Carbohydrate Active Enzymes
LCB: lignocellulosic biomass
GH: glycoside hydrolases
GT: glycosyl transferases
PL: polysaccharide lyases
CE: carbohydrate esterases
AA: auxiliary activities
CBM: carbohydrate-binding modules

## Ethics approval and consent to participate

Not Applicable

## Consent for publication

Not Applicable

## Availability of data and materials

Data used in this analysis has been procured from open-source databases such as NCBI. Raw and analyzed data along with scripts can be provided upon request.

## Authors’ contributions

GS generated the idea and provided guidance. NS performed the analysis and wrote the first manuscript text. NS and GS prepared Figures and Tables. All authors reviewed the manuscript.

## Acknowledgments

Not Applicable

